# Actinidia seed-born latent virus is transmitted paternally and maternally at high rates

**DOI:** 10.1101/2021.02.19.432071

**Authors:** NT Amponsah, R van den Brink, PM Datson, PT Austin, M Horner, RM MacDiarmid

## Abstract

Actinidia seed-borne latent virus (ASbLV, Betaflexiviridae), was detected at high frequency in healthy seedlings grown from lines of imported seed in a New Zealand post-entry quarantine facility. To better understand how to manage this virus in a dioecious crop species, we developed a rapid molecular protocol to detect infected progeny and to identify a reliable plant tissue appropriate to detect transmission rates from paternal and maternal parents under quarantine environment.

The frequency of ASbLV detection from true infection of F1 progeny was distinguished by testing whole seeds and progeny seedling tissues from a controlled cross between two unrelated parents; an ASbLV-infected staminate (male) plant and an uninfected pistillate (female) plant, and the process was repeated with an ASbLV uninfected staminate (male) plant and an infected pistillate (female) plant. Individual whole seeds, or single cotyledons from newly-emerged seedlings, true leaf or a root from those positive-tested seedlings, were assessed for presence of ASbLV by reverse transcription-polymerase chain reaction (RT-PCR) analysis. The virus was detected at a high incidence (98%) in individual seeds, but at a much lower incidence in seedling cotyledons (62%). Since detection results were consistent (*P*=95%) across the three seedling tissues (i.e. cotyledons, leaves and roots) only cotyledons were tested thereafter to determine ASbLV transmission to F1 progeny. F1 seedlings from three crosses were used to compare transmission rates from infected staminate versus infected pistillate parents. One cross from a single flower used an uninfected pistillate vine pollinated by an infected staminate vine, and two crosses (also from a single flower) used an infected pistillate vine (a sibling of the infected staminate vine), pollinated by either of two unrelated uninfected staminate vines.

Cotyledon testing of seedlings from each cross confirmed staminate transmission at high frequency (∼60%), and pistillate transmission at even higher frequency (81% and 86%, respectively).

The results show ASbLV is transmitted at very high rates, whether from infected ovules or pollen. Transmission to seedlings is lower than detection in whole seeds perhaps due to ASbLV being sometimes residing on (or within) the seed coat only. The results also show RT-PCR of cotyledons allows non-destructive detection of ASbLV in very young seedlings, and could be used to screen kiwifruit plants in a nursery to avoid virus spread to orchards. Likewise, bulk testing of seed lots can quickly detect infected parent plants (fruit bearing female or male pollinator) already in an orchard.

**Importance:** Actinidia seed-borne latent virus (ASbLV, Betaflexiviridae), was detected at high frequency in healthy seedlings grown from lines of imported seed in a New Zealand post-entry quarantine facility. However there are several technical barriers to detecting the presence of seed transmitted viruses and understanding their biology, which has significance for detection in quarantine and subsequent management under germplasm collections. To overcome this, we developed a rapid molecular protocol to detect infected progeny and to identify a reliable plant tissue appropriate to detect transmission rates from paternal and maternal parents under quarantine environment. Individual whole seeds, or single cotyledons from newly-emerged seedlings, true leaf or a root from those positive-tested seedlings, were assessed for presence of ASbLV by reverse transcription-polymerase chain reaction (RT-PCR) analysis. This was done with seed lots obtained from four separate controlled crosses between ASbLV-infected and ASbLV-uninfected Actinidia chinensis var. deliciosa parents.

## 1 Introduction

Intergenerational virus transmission via seed offers a narrowly-targeted mode of transmission to successive generations of genetically-similar plant hosts that might be expected to share similar susceptibility to infection as the seed’s parents. This contrasts with mechanical or vector transmission of virus infection, which allows distribution beyond the current host’s gene pool. Seed transmission can allow successive generations of infection, replication and spread of the virus as seeds are distributed according to the host’s reproductive and ecological strategy. It may also provide an on-going opportunity for a mutualistic relationship, inter-generationally, between plant host and genetic “guest” (Johansen et al. 1996). However, many viruses has been described to be seed transmitted and that at least 25% of plant viruses are vertically transmitted (Simmons and Munkvold 2014).

Seed infection or the virus present in or on seed may result from either the paternal or maternal parent, or both (Albrechtsen 2006). Maternal transmission can result from: indirect invasion of the embryo via infected meristematic tissue and derived megaspore mother cells; direct invasion of the embryo via the suspensor as a transient pathway; or by infection of maternal seed parts (e.g., integuments of the ovule) without embryo invasion. For instance, the Potyvirus Pea seed-borne mosaic virus is able to infect the maternal tissue of the micropylar region, move via symplastic pore-like openings between the testa and endoplasm, to access the suspensor cells and directly invade the embryo in early development (Roberts et al. 2003). Paternal transmission can result from viruses being carried on or in pollen tubes to infect the embryo or endosperm at fertilisation (Isogai et al. 2015) or by viruses escaping from pollen tubes and thence onto or into the ovule (i.e., the prospective seed). Furthermore, even if the embryo remains virus-free, the process of germination can result in mechanical infection of the emerging seedling via physical interaction with a virus infested seed coat. For example, Ribgrass mosaic virus (RMV), a Tobamovirus that infects *Actinidia* (Chavan et al. 2012), can be transmitted from the surface of the seed coat.

There are several technical barriers to detecting the presence of seed transmitted viruses and understanding their biology, which has significance for detection in quarantine and subsequent management under germplasm collections. Firstly, the seed transmission mode favours viruses that are not deleterious to their host and thus generally form asymptomatic infections (Villamor et al. 2019), which are not easily recognised and are therefore under-represented within the literature. This mode of transmission has likely been a significant contributor to the unprecedented number of asymptomatic viruses identified by high throughput sequencing over the last decade (Villamor et al. 2019).

Limited knowledge of seed-transmission in the context of plant material movement increases uncertainty associated with the biosecurity risk of inadvertent virus movement and transmission. In general, the import of seeds or pollen is regarded as having lower risk of pest and pathogen contamination than other planting material, but transmission of seed-borne viruses remains a risk to be managed (Maule et al 1996; Card et al. 2007).

The first *Actinidia* seed were introduced to New Zealand in 1904, and produced the first fruit outside China 6 years later (Smith 2000). Since then, kiwifruit has grown to become an important crop in New Zealand, with development of new cultivars supporting worldwide marketing and production. Virus-like symptoms had been reported in kiwifruit (Caciagli and Lovisolo 1987) but no viruses were identified until 2001, when a strain of Apple stem grooving virus (ASGV) was detected in post-entry quarantine, in plants of *Actinidia chinensis* imported from China (Clover et al. 2003). Since then, a total of 16 viruses, classified into three groups, have been found in kiwifruit growing areas including China, Italy and New Zealand, including Ribgrass mosaic virus (RMV), first detected in *Actinidia chinensis* var. *deliciosa* and var. *chinensis* held in post-entry quarantine in New Zealand (Chavan et al. 2012; Chavan et al. 2013; Chavan and Pearson 2016; Zhao et al. 2019; Zhao et al. 2020). The list of viruses identified in *Actinidia* continues to grow (Blouin et al. 2013;; Zheng et al. 2014; Biccheri 2015; Lebas et al. 2016; Zheng et al. 2017; Blouin et al. 2018; Veerakone et al. 2018; Wang et al. 2018; Wen et al. 2018; Zhao et al. 2019; Wang et al. 2020; Zhao et al. 2020), including detection of viruses first identified in other species (James and Phelan 2016).

Veerakone et al. (2018) described the assembly and analysis of the genome of Actinidia seed-borne latent virus (ASbLV), a member of the Betaflexiviridae, from *Actinidia* plants in New Zealand *Actinidia* species, commercialised as ‘kiwifruit’ and are bred from wild germplasm sourced from China, the centre of biodiversity for the genus. The virus was also identified in imported seed within a New Zealand post-entry quarantine facility and was subsequently identified in related and healthy-appearing *Actinidia* seedling collections (Veerakone et al. 2018). Tracing of the imported *Actinidia* families identified some seedlings growing in germplasm collections within New Zealand, and revealed a high seed detection rate (98%).

The New Zealand kiwifruit industry mainly uses seed propagated rootstocks with clonally propagated and grafted scions; there are also some clonally propagated rootstocks available. The majority of kiwifruit material that has come through quarantine in New Zealand has been seed, although both pollen and budwood have also been imported. Since *Actinidia* species are dioecious, viruses may be transmitted via pollen from the male (staminate) plants and/or from the female (pistillate) plants that bear the ovaries that form the fruit. Ovules in each fruit are collectively fertilised by approximately a thousand pollen grains, with each successful pollen grain resulting in a seed within a single fruit. In a quarantine context, asymptomatic infections undermine the importance of inspection methods, and unknown prevalence and infection rates make designing cost-effective sampling regimes more difficult, especially as seed testing is typically destructive.

Limited knowledge of presence, and infection incidence, of seed- and pollen-transmitted viruses is also significant in the context of germplasm management. In plant breeding viruses may be vertically or horizontally transmitted between generations. However, virus-free progeny are generally preferred because viruses may prove detrimental in new host genotypes. Other infections if present may lead to synergistic symptom expression. The aim of this study was to determine a process to identify a reliable tissue to detect transmission of ASbLV to F1 progeny under a quarantine environment including both (a) a rapid molecular method, and (b) the most reliable tissue to use for testing.

## 2 Materials and methods

To compare ASbLV transmission rates from micro- and mega-gametophytes we crossed unrelated, uninfected sexual partners with an infected partner. To study transmission from the mega-gametophytes, the same infected female was crossed with two unrelated males. For transmission from the micro-gametophyte, two unrelated, infected males were crossed with two unrelated females; the infected female and one of the infected males were half-siblings, having originally been derived from seeds of the same fruit. Binomial significance test was performed on ASbLV transmission data.

### 2.1 Seed material

Seed lots ‘C15’, ‘C53’, ‘T66’ and ‘X84’ were obtained as single-fruit extracted seed sublots derived from four separate controlled crosses performed between ASbLV-infected and ASbLV-uninfected *Actinidia chinensis* var. *deliciosa* parents (Figure 1). Both the infected and non-infected parents were originally sourced from seed derived from open-pollenated fruit collected from wild vines.

**Figure 1.**
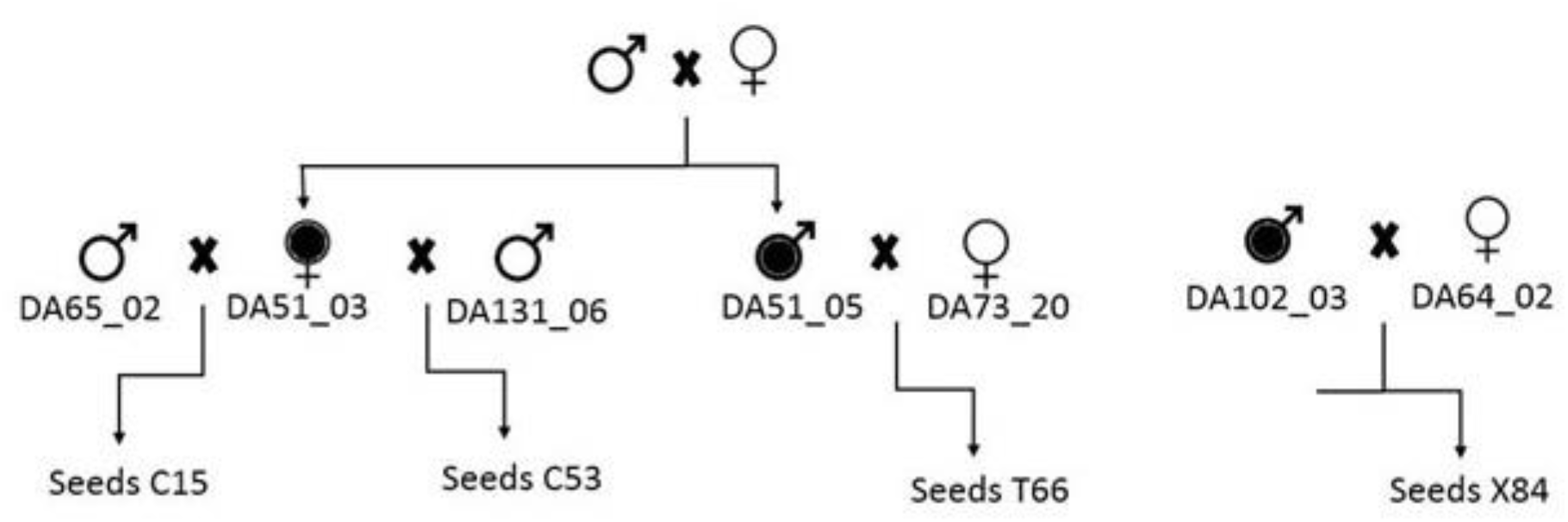
Identification and relationship of parents used for crosses that generated seeds tested for *Actinidia latent seed-borne virus* (ASbLV) transmission from maternal and paternal *Actinidia deliciosa* parents. Seeds of lot ‘X84’ were tested destructively as whole seeds, or grown before cotyledons, true leaves and roots were tested. Seeds of lot ‘C15’, ‘C53’ and ‘T66’ were tested by assay of cotyledons only. Open circles indicate uninfected parents and filled circles indicate ASbLV infected parents

A range of crosses between ASbLV-infected and non-infected parents were performed in December 2015 using field-grown vines. The quarantine facility housed uninfected Actinidia plants grown under a level three facility which were used as negative controls. These plants had previously been tested by RT-PCR as ASbLV negative. The ASbLV infection status of the parents was assessed using reverse transcription-polymerase chain reaction (RT-PCR) targeting a 278 bp sequence of the RNA-dependent RNA polymerase (RdRP) gene of the virus by testing leaf samples collected in March and December 2015 from field-grown vines and repeated in August 2016 by testing newly shoot buds just emerging from dormant budsticks. (Details are provided section 2.3). On female vines the terminal ends of shoots containing flower buds were covered with a paper bag prior to anthesis then sealed by folding and stapling so that the flowers were completely enclosed in the bag. Flower buds from male vines were collected just before the flower buds opened, anthers were extracted from the flower buds and air-dried to extract the pollen. Pollinations were then performed in the field by briefly opening the paper bags once the flowers inside had opened, and applying the isolated pollen using a small paint brush, Different brushes were used for each pollination. The flowers were then sealed in the paper bags again and left until fruitlets formed (December) when each paper bag was exchanged with an onion bag (5 mm mesh bag). Fruit from the crosses were harvested in May 2016 and seeds were extracted.

Whole seeds from lots ‘X84’, ‘T66’, ‘C15’ and ‘C53’ extracted from single fruit, respectively were tested for the presence of ASbLV (Table 1). Each seed lot consisted of over 1000 seeds extracted from a single kiwifruit berry. Seed from lots ‘C15’ and ‘C53’ are from the same ASbLV-infected *Actinidia* mother plant (DA51_03), which is a sibling to the infected male DA51_05 used to produce seed lot ‘T66’ (Figure 1). Seed lot ‘X84’ was generated using an unrelated infected male DA102_03.

**Table 1:** Identification and relationship of parents used for crosses that generated seeds tested for *Actinidia latent seed-borne virus* (ASbLV) infection. Seeds of lot ‘X84’ were tested destructively as whole seeds, or grown before cotyledons, true leaves and roots were tested. Seeds of lot ‘C15’, ‘C53’ and ‘T66’ were tested by assay of cotyledons only

| Cross ID | Pistillate vine<br>ID ♀ | Staminate vine<br>ID ♂ | Total samples (n) | Total infected | ASbLV detection rate (%) | Confidence intervals |
| --- | --- | --- | --- | --- | --- | --- |
| ‘X84’<br>Whole seed | DA64_02<br>(uninfected) | DA102_03<br>(infected) | 50 | 49 | 98 | 89-100 |
| ‘X84’<br>cotyledons | DA64_02<br>(uninfected) | DA102_03<br>(infected) | 50 | 32 | 64 | 47-75 |
| ‘X84’<br>Juvenile<br>(leaf) | DA64_02<br>(uninfected) | DA102_03<br>(infected) | 32 | 18 | 56 | 39-75 |
| ‘X84’<br>Juvenile<br>(Roots) | DA64_02<br>(uninfected) | DA102_03<br>(infected) | 20 | 12 | 60 | 36-81 |
| ‘X84’<br>Young<br>leaves | DA64_02<br>(uninfected) | DA102_03<br>(infected) | 13 | 8 | 61 | 32-86 |
| ‘X84’<br>Mature<br>leaves | DA64_02<br>(uninfected) | DA102_03<br>(infected) | 13 | 8 | 61 | 32-86 |
| ‘C15’<br>(Cotyledons<br>) | DA51_03<br>(infected) | DA65_02<br>(uninfected) | 53 | 43 | 81 | 68-91 |
| ‘C53’<br>(Cotyledons<br>) | DA51_03<br>(infected) | DA131_06<br>(uninfected) | 46 | 35 | 76 | 61-87 |
| ‘T66’<br>(Cotyledons<br>) | DA73_20<br>(uninfected) | DA51_05<br>(infected) | 49 | 29 | 59 | 44-73 |

NB: DA51\_05 is a sibling to pistillate parent DA51\_03

### 2.2 Isolation of total RNA from seeds, cotyledons, leaves and roots

To test for the presence of ASbLV, total RNA was isolated from seeds, cotyledons, leaves (young and mature) and roots from parental and/or progeny *A. chinensis* (details provided in section 2.3). For reasonable grounded seed material for processing, each RNA extraction used one test seed that was combined with four *Actinidia* seeds from another source known to be non-infected with ASbLV. For the extraction of total RNA from cotyledons, a single cotyledon weighing (1 to 8 mg depending on age) was harvested from each seedling 1–2 weeks after emergence. For total RNA extraction from leaf tissue, a fully formed and opened leaf (∼10 mg) was harvested from a seedling 3–4 weeks after seedling emergence. For root tissue, a piece of root tip (∼100 mg) was harvested from a seedling 3–4 weeks after emergence. Tissue isolated from plants that had previously been tested by RT-PCR as ASbLV negative and grown under level 3 facility) or plants tested by RT-PCR and known to be infected with ASbLV (grown within a PFR experimental orchard in Motueka, New Zealand) were used as negative and positive controls, respectively.

A silica milk-based method developed by Menzel et al. (2002) was used for RNA extraction. Extraction buffer (6 M guanidine hydrochloride, 0.2 M sodium acetate pH 5.0, 25 mM EDTA, and 2.5% PVP-40) and the plant sample was ground within a grinding bag (10 cm × 8 cm, BIOREBA AG™, Switzerland) using a ball-bearing grinder. The ground extract (750 µL) was aliquoted into a new Eppendorf tube to which 100 µl SDS (10%) was added and the mixture incubated at 70°C for 10 min with intermittent shaking, incubated on ice for 5 min, then centrifuged at 15,000 × g for 10 min to remove the solid material. The supernatant (300 µL) was transferred to a new tube and 300 µL sodium-iodide-solution (6 M, stabilized by 0.15 M sodium sulfite), 150 µL ethanol (99.6%) and 25 µL silica-suspension (1g/ml silicon dioxide, Sigma-Aldrich S5631, pH 2) were added. The contents were shaken continuously for 10 min then the mixture was centrifuged at 2000 × g for 1 min and the pellet washed twice by re-suspending the pellet in 500 μL wash buffer (10 mM Tris-HCl pH 7.5, 0.05 mM EDTA, 50 mM NaCl and 50% ethanol). After washing and air drying the pellet, the RNA was released from the dry silicon dioxide pellet by dissolving in 100 μL Tris EDTA buffer (10mM Tris-HCl pH 7.5, 0.05 mM EDTA) and incubating at 70°C for 4 min, to release the nucleic acids. The final supernatant was centrifuged at 2000 × g for 5 min and transferred to new tubes and stored in a freezer at -20°C prior to RT-PCR analysis.

### 2.3 Virus detection by RT-PCR

ASbLV was detected by one-step RT-PCR and RNA quality was assessed by amplification of the plant NAD5 using specific primers designed in this study, i.e. BetaRDRP-F2: GAATCAGACTATGAAGCATTTGATGC and BetaRDRP-R2: CACATATCGTCACCTGCAAATGCTATTG, that amplified a 278 bp product from the ASbLV RdRp. Nad5 mRNA-specific primers Nad5-F and Nad5-R (mRNA coding mitochondrial gene of higher plants encoding subunit 5 of the NADH ubiquinone oxidoreductase complex) described by Menzel et al. (2002) was used as an internal control. Each reaction contained PCR 10x buffer (Invitrogen™, ThermoFisher Scientific, USA), DEPC-treated water, 50 mM MgCl_2_, 10 mM dNTPs, 60 µM primers, 100 mM DTT, 0.5U Platinum^®^ Taq DNA polymerase and 10U SuperScript™ III reverse transcriptase (ThermoFisher Scientific, USA). Amplification using a thermal cycler (Eppendorf MasterCycler Gradient, Netheler-Hinz GmbH, Hamburg) was achieved by the following temperature regime: Reverse transcription, 30 min for 48°C; Taq activation, 3 min at 96°C; followed by 35 cycles of amplification 30 s at 95°C, 30 s at 56°C and 45 s at 72°C, with a final extension of 2 min at 72°C. Amplicons were separated by electrophoresis in a 1.5% Ultrapure™ agarose gel in 0.5 × TBE buffer at 10 V/cm for 50 min. The gel was stained with 0.5 μg/ml ethidium bromide for 15 min, and visualised with ultraviolet light.

### 2.4 Seedling growth

Prior to sowing, 100 *Actinidia* seeds from each test cross were surfaced sterilised in 1% bleach (Sodium hypochlorite) and air dried under laminar flow prior to soaking in 1 mL of 3000 ppm gibberellic acid for 24 h, then spread over 100 mm diameter moist Whatman^®^ filter paper (Buckinghamshire, United Kingdom). Each seeded filter paper was placed on top of a half filled pasteurised medium grade bark/pumice potting media (Daltons Ltd Matamata) in 150 mm punnet container and covered with 4 mm particle size sterile sand. Seeds were germinated at 22.5°C for 1 week and 50 healthy looking seedlings were individually transplanted into 55 mm plastic pots filled with potting mix for growth. Subsequent tests for ASbLV transmission in cotyledons, leaves and roots were carried out when the plants were 1-4 weeks old (as detailed in section 2.2).

The ratios from male and female sources of infection, and the ratios from the different tissue types of the ‘X84’ seed lot were compared using binomial generalized linear models.

## 3. Results

Successful amplification of the NAD5 reference part of an RNA molecule by one-step RT-PCR showed that all RNA extracts from whole seeds, roots, leaves and cotyledons were competent for RT-PCR after a PCR check showing no DNA in the RNA preparation. ASbLV RNA from infected *Actinidia* seeds, roots, leaves and cotyledons was successfully amplified by RT-PCR. Negative controls showed bands for NAD5 only, and blank extractions (buffer or water controls) did not amplify (Figure 2).

**Figure 2.**
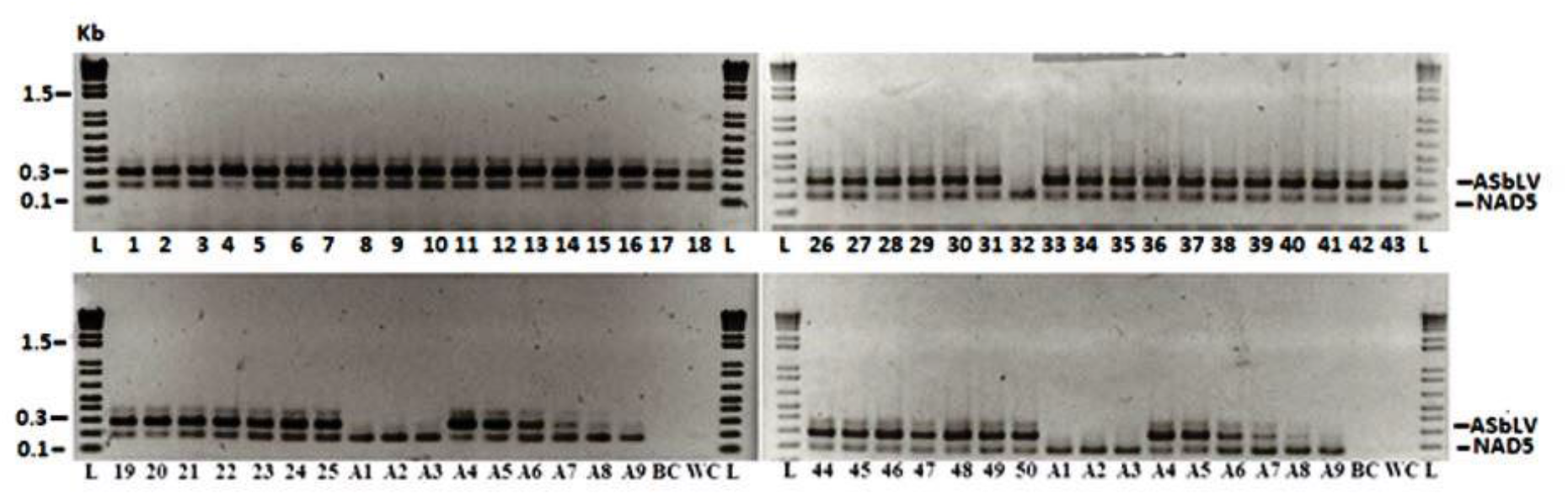
*Actinidia latent seed-borne virus* (ASbLV) reverse transcription-polymerase chain reaction (RT-PCR) product showing the performance of the primer pairs on positive and negative samples. Lanes L is 1kb+ ladder (Invitrogen™); the numerical numbered lanes 1-50 represents the test samples; Lanes A1–A3 are *Actinidia* seed negative controls; Lanes A4–A9 are positive control dilution series (10^0^ to10^−5^); BC buffer control; WC water control. ASbLV is amplicon size is 278bp and NAD5 internal control is 181bp

### 3.1 Whole seed testing

An ASbLV amplification band matching the expected size of 278 bp (Figure 2) was amplified from all except one whole seed (lane 32) from *Actinidia* seed lot ‘X84’, along with the band for the NAD5 internal RT-PCR control (181bp). This ASbLV detection rate (*n* = 49/50) provides a binomial confidence interval of 89–100% infection, *P*=95% (Table 1).

### 3.2 Transmission from infected staminate parents

ASbLV was detected in approximately two-thirds of germinated seedlings from the same ‘X84’ seed lot (*n*=32/50 detected; confidence interval 47–75% infected, *P*=95%), when a cotyledon from each seedling was tested individually (Table1). When juvenile leaves produced from the 32 infected seedlings (positive by cotyledon assay) were tested, approximately half of those leaves tested were positive (Table 1) for the ASbLV (*n*=18/32; confidence interval 39–75% infected, *P*=95%). Of the 32 infected plants, only 13 had multiple fully formed leaves after 2 weeks’ growth. When two leaves from the same plant (one younger and one older) were tested from each of these 13 seedlings, about half of the plants (for both leaf ages) tested positive for ASbLV (*n*=8/13; confidence interval 32–86% infected, *P*=95%) (Table 1).

Roots sampled from 20 surviving ASbLV-positive seedlings (the 13 positive seedlings and an additional seven with no fully formed leaves whose leaves and/or cotyledons, respectively, had previously tested ASbLV-positive) were tested for the virus, a little over half of that number (Table 1) tested positive for ASbLV (*n*=12/20; confidence interval 36-81% infected, *P*=95%). ASbLV detection in seedling leaves and roots matched that observed for each respective cotyledon but was lower than the detection frequency for lot ‘X84’ whole seeds.

ASbLV was detected in an equivalent proportion of seedlings from the ‘T66’ seed family (resulting from a cross of an infected *Actinidia* staminate parent DA51_05, unrelated to DA102_03, with a non-infected pistillate parent DA73_20), when cotyledons were individually tested by RT-PCR (*n*=29/49; confidence interval 44-73% infected, *P*=95%). This detection rate indicates transmission of ASbLV from the infected staminate parent to about two thirds of ‘T66’ seedlings is similar to the rate of transmission observed for the ‘X84’ seed family.

### 3.3 Transmission from infected pistillate parents

ASbLV was detected in a high proportion of cotyledons from *Actinidia* seedlings grown from two seed families with an infected pistillate parent. ASbLV was detected in 81% of seedling cotyledons grown from seed lot ‘C15’ (seed from crossing infected pistillate plant DA51_03, a sibling to infected plant DA51_05, with the unrelated non-infected staminate plant DA65_02, Fig. 1) when tested by RT-PCR (*n*=43/53; confidence interval 68-91% infected, *P*=95%). The same result was observed for a second set of germinated *Actinidia* seedlings, family ‘C53’ (seed from crossing the same infected pistillate plant DA51_03 with the non-infected staminate plant DA131_06, not known to be related to any other plant in this study, Figure 1). In this case, ASbLV was detected in 85% of cotyledons sampled from ‘C53’ seedlings (*n*=35/46; confidence interval 61-87% infected, *P*=95%) when individually tested for ASbLV (Table 1). The results indicate a very high rate of transmission of ASbLV from infected pistillate parents, on average possibly slightly higher than the transmission rate from infected staminate parents. Comparison of ASbLV detection in cotyledons from crosses involving maternal- and paternal-infection showed a significantly higher proportion of infected seedlings arising from maternal infection (deviance = 7.8, 1 df, *P*=0.005). There was no significant difference in transmission frequency of ASbLV to different seedling tissues in the case of the one infected paternal parent crossing (‘X84’, Table 2) where this was investigated (deviance = 0.1, 2 df, *P*= 0.967).

**Table 2.**
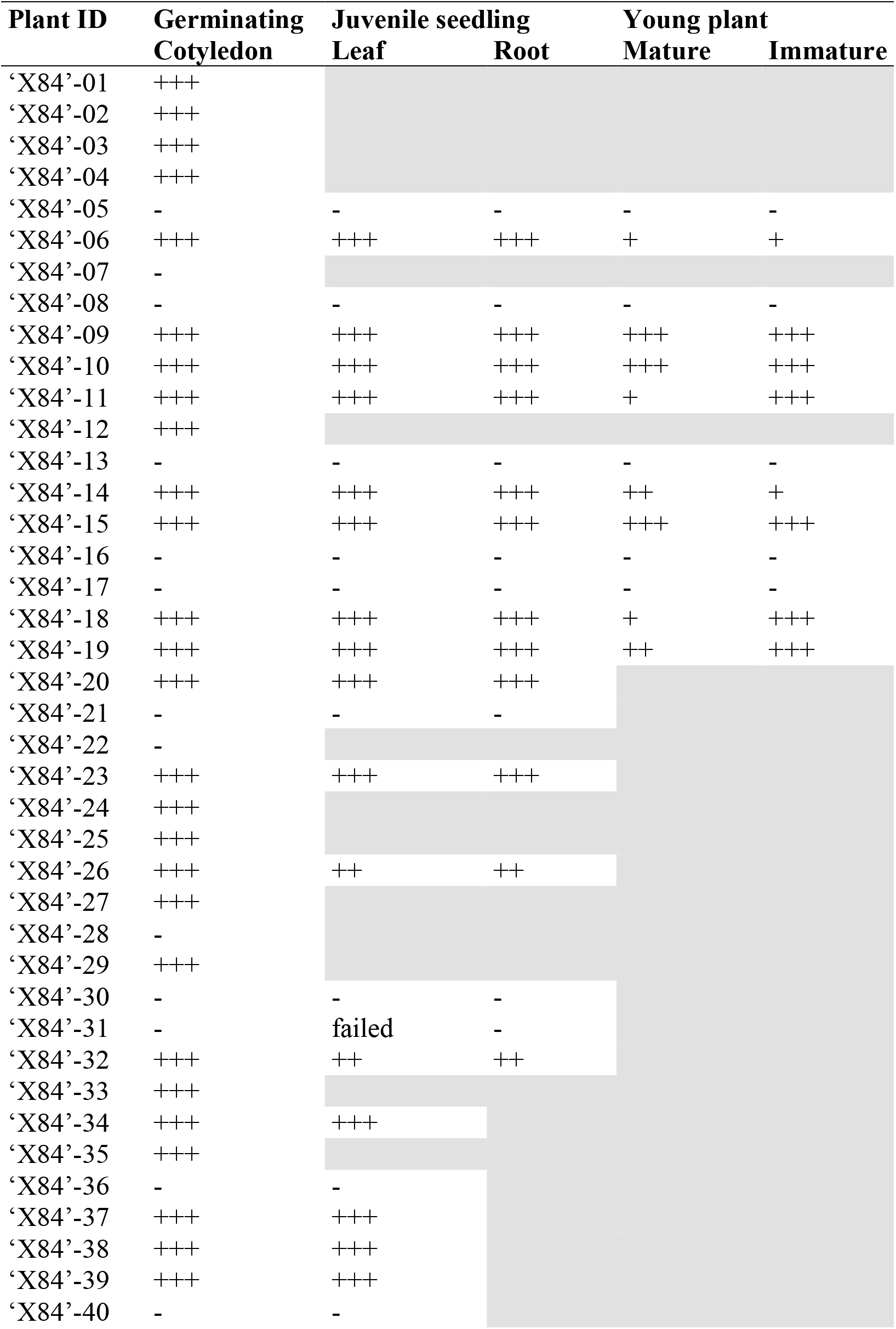
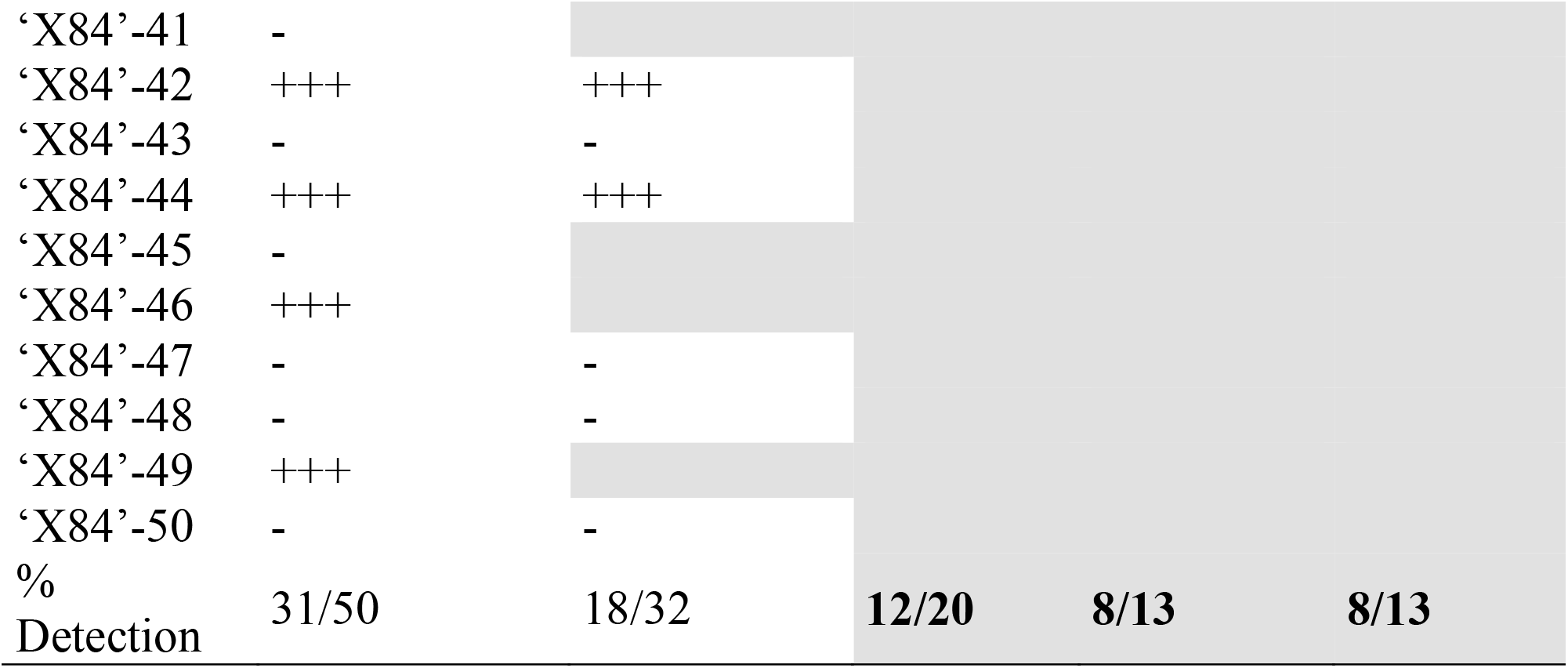
Infection status of *Actinidia* seedlings from ‘X84’ cross, tested by end-point reverse transcription-polymerase chain reaction (RT-PCR) for RNA of *Actinidia seed-borne latent virus* and using RNA for the plant NAD5 gene as an internal reference. Grey backgrounds indicate dead plants. (PCR target band strength: +++ = much stronger than NAD5 reference; ++ = similar to NAD5 reference, + = weaker than NAD5 reference, – = target not detected.)

## 4 Discussion

One of the major factors contributing to plant virus long-distance dispersal is the global trade of seeds. In this study, we discovered that the transmission frequency of ASbLV to seedling tissues from maternal or paternal parents differed significantly depending on the floral function of infection source (ovule or pollen). The frequency of infection was highest in the seeds demonstrating the virus is seed borne either on or within the seed. For virus transmission from the infected seeds to seedlings, the infection in the cotyledons was higher than in the juvenile or matured leaves. This may reflect the interplay between the plant’s growth, the viral infectious cycle and the plant’s defence responses that result in virus replication being slowed allowing the plant growth to progress faster than virus transmission through the nascent organs. This pattern mimics those described by Domier et al. (2011) whereby plant defence responses regulate virus virulence by altering the virus distribution in the plant. Furthermore, in our transmission study more cotyledon samples than mature or juvenile leaves were tested due to the attrition following the cotyledon sampling process, despite careful handling. The removal of a cotyledon may have significantly weakened several of the seedlings.

Because ASbLV has not been associated with host symptoms, it is presently assessed within New Zealand as low-to-zero biosecurity risk. Expanding awareness of the multiple viruses found in *Actinidia* is important for commercial fruit production, especially in an export-oriented industry context such as the kiwifruit industry. Breeding now involves a wide range of genotypes and regional sources for inter-varietal or inter-specific fertilisation to produce parental lines and seedlings for elite cultivar selection. Given the novelty of this crop, this large-scale intensive crossing programme creates a significant risk of introducing unrecognised viruses into breeding programmes. While latent in the parents, it may be possible that the the virus transmission could create novel disease in the new host genotype, or on expansion of the industry to new environments (such as indoor production), which may be conducive to damaging symptom expression.

The relatively high frequency of ASbLV transmission to progeny found in this study raises a number of practical management and scientific research questions. Many members of the Betaflexiviridae show latency and are symptomatic only in specific situations, e.g. associated with graft incompatibility. It may also be likely ASbLV co-evolved with the wild *Actinidia* species and its presence in multiple wild-sourced accessions suggests a possible benefit to the host, supporting its persistence in the host plant and further research needs to be done to confirm or deny that. For example viruses such as Brome mosaic virus, family Bromoviridae, Cucumber mosaic virus, family Bromoviridae, Tobacco rattle virus, family Virgaviridae, and Tobacco mosaic virus (TMV), family Virgaviridae are known to be beneficial to crops because they are known to confer tolerance to drought and freezing temperatures in several different crops (Roossinck, 2011). Surveys in China, or of China-derived seed, may reveal sequence variation within or even between *Actinidia* species that will support their co-existence. Empirical biological testing of ASbLV infected and uninfected genotypes needs to be performed to determine any benefit to the host under abiotic or abiotic stress. For instance, it would be intriguing to determine whether the virus provides any benefit to kiwifruit challenged with *Pseudomonas syringae* pv. *actinidiae* (Psa).

In New Zealand, imported breeding and propagation material, such as seedlings grown from seed, not only undergoes government-specified testing and inspection in post-entry quarantine but also industry-agreed testing for disease pathogens prior to use in breeding or propagation.

Crosses using an ASbLV-infected maternal parent averaged ∼80% transmission (total of 78 from 99 seedlings), while crosses using an ASbLV-infected paternal parent averaged ∼60% transmission (total of 60 from 99 seedlings). If in planta movement of ASbLV is passive via cell division only, this may offer an interpretation for the possibility, based on results from the limited number of crosses studied, that transmission frequency from the paternal parent is lower than from the maternal parent. If the suggested lower transmission rate via pollen is representative of other crosses, then it is possible this may reflect loss of the passively-distributed virus within the fast-growing pollen tube of the microgametophyte. This could be a result of simple dilution of the virus in the pollen tube, and passive exclusion from some germ cells at the tube tip, or it could reflect a more active process of virus exclusion by the microgametophyte. Passive movement is a hallmark of persistent viruses that like ASbLV do not display disease symptoms, but unlike ASbLV lack a movement protein yet are present in all host cells (Roossinck 2010). The dioecious nature of Actinidia may provide a biological lever to prise apart cell division and ASbLV movement to understand its transmission biology.

While the scope of this initial study is not sufficient to offer a conclusive interpretation of the suggested difference between maternal and paternal transmission, the possibility has attractive implications for practical management and use of infected parental populations. As identified here, 36% seeds from a cross involving only a single ASbLV infected parent yielded seedlings without detectable virus even though the virus may be carried on or within the seed coat.

The high and directionally-balanced transmission of ASbLV (i.e., seed infection from both the maternal and paternal sporophyte parents), along with absence of symptoms is consistent with the virus being well-habilitated as a ‘guest’ within *Actinidia* as a host species. Even where infection is asymptomatic, virus infection of new cultivars is undesirable due to perceived risks that may limit opportunities for their commercial exploitation. According to Cobos et al. (2019) more than 25% of plant viruses can infect seeds, and start new infections in areas previously not present, however, the infection traits associated with the efficiency of virus seed transmission are largely unknown. The frequency of infection at 60–80% is not exceptionally high for seed-borne viruses. For instance, it is within the transmission frequency range of Pea seed-borne mosiac virus into susceptible *Pisum sativum* genotypes (Maule and Wang 1996). As indicated by Alizon et al. (2009), a pathogens ability to be transmitted is arguably the most important determinant of its parasitic fitness and most theoretical models of the evolution of parasites consider infection traits, such as virulence, as relevant factors for parasite fitness because they affect the efficiency of between-host transmission. The near-even reciprocity is similar to seed infection of *Arabidopsis thaliana* by Turnip yellow mosaic virus (TYMV), where virus from either the female or the male parent could invade the seed, at high frequency although it contrasts with the relationship of *A. thaliana* and TMV, for which the maternal parent was the only route by which TMV invaded the seed (de Assis Filho and Sherwood 2000).

To date there is also no evidence for mechanical transmission of ASbLV to indicator species including *Chenopodium quinoa, C. amaranticolor, Cucumis sativus, Nicotiana glutinosa, N. benthamiana, N. tabacum* (var. Samsun), *N. occidentalis* 37B, *N. clevelandii*, and *Phaseolus vulgaris* var. Thus, if the virus is solely transmitted vertically via the micro- and megagametophyte to the embryo and thence to the seedling, it is plausible that ASbLV moves only passively via cell division to the subsequent daughter cells that eventually form the mature sporophyte. Such a transmission mode via cell division of the host plant would not require either cell-to-cell or long distance trafficking of the virus; indeed movement proteins would be redundant (Veerakone et al. 2018).

Cytological studies of ASbLV distribution and movement during fertilization, in the embryo and in grafted plants (e.g., reciprocal grafting of infected and uninfected stocks and scions, or inter stocks), may allow elucidation of the mode of transmission of ASbLV within its host. In particular, given the possibility of virus dilution or exclusion in the pollen tubes, an intriguing possibility is that transmission frequency may be related to pollen tube length. In which case, seeds at the proximal (stalk) end of fruit from crosses using infected paternal parents could offer lower frequencies of virus transmission compared to seeds at the distal (stylar) end, allowing more efficient recovery on non-infected seedlings. This possibility needs to be explored in further reciprocal crossing studies with ASbLV.

## 5 Acknowledgement

We thank the following people for their practical help and encouragement: Alison Duffy (The New Zealand Institute for Plant and Food Research Limited, PFR), PEQ project support; Duncan Hedderley (PFR), statistical analysis; Guy Middlemiss (PFR), site facilities management; plant propagation; Nicola Mauchline (PFR), plant materials tracking; Shirley Day, Jaswinder Sekhon, Thomas Paterson, Natalie Taufa and Kirsten Hoeata (all of PFR, Te Puke) for cross pollination, fruit harvests and seed extraction and vine management; Lisa Ward (MPI), diagnostics liaison; Michael Tuffin (Mantis Greenhouse Solutions).

